# Method for modeling oviduct function and impact on embryonic development

**DOI:** 10.64898/2026.08.06.743297

**Authors:** Kalli K Stephens, Vakil Ahmad, Maria A. Silva, Makenna K Shifflett, Jiude Mao, Jason A. Rizo, Mark I Hunter, Andrew M Kelleher, Wipawee Winuthayanon

**Affiliations:** Division of Animal Sciences, College of Agriculture, Food and Natural Resources, University of Missouri, Columbia, Missouri, USA 65211; Department of Obstetrics, Gynecology, and Women’s Health, University of Missouri, Columbia, Missouri, USA 65211; Department of Medical Pharmacology and Physiology, School of Medicine, University of Missouri, Columbia, Missouri, USA 65211

## Abstract

Direct experimental analysis of the mammalian oviduct is constrained by limited tissue access and the short lifespan of *ex vivo* preparations. Extracellular matrix-embedded three-dimensional epithelial organoids provide longer-term *in vitro* models. However, their inward-facing apical surface and the absence of supporting stromal cells limit physiological studies of the oviduct, including ciliary activity and maternal-embryonic interactions. Here, we provide a step-wise protocol detailing the generation of mouse and human oviductal assembloids in which epithelial cells form an outward-facing (apical-out) layer around a stromal core. Epithelial and stromal cells from adult mouse oviducts or human Fallopian tubes are isolated, expanded separately, and subsequently aggregated in a rotational culture system. The protocol also outlines morphological and immunostaining criteria for confirming cellular organization, whole-mount detection of external cilia, measurement of ciliary beat frequency, and co-culture of mouse assembloids with preimplantation embryos. Mouse and human assembloids retained epithelial and stromal identity and displayed cilia at the accessible outer surface. In a proof-of-concept experiment, embryos co-cultured with the assembloids developed to blastocysts at a rate similar to that of *in vivo*-derived blastocysts. This reductionist system provides a straightforward and tractable model to investigate oviduct physiology and embryo-maternal communication while allowing direct manipulation and observation of the epithelial interface.

**Graphical Abstract:** 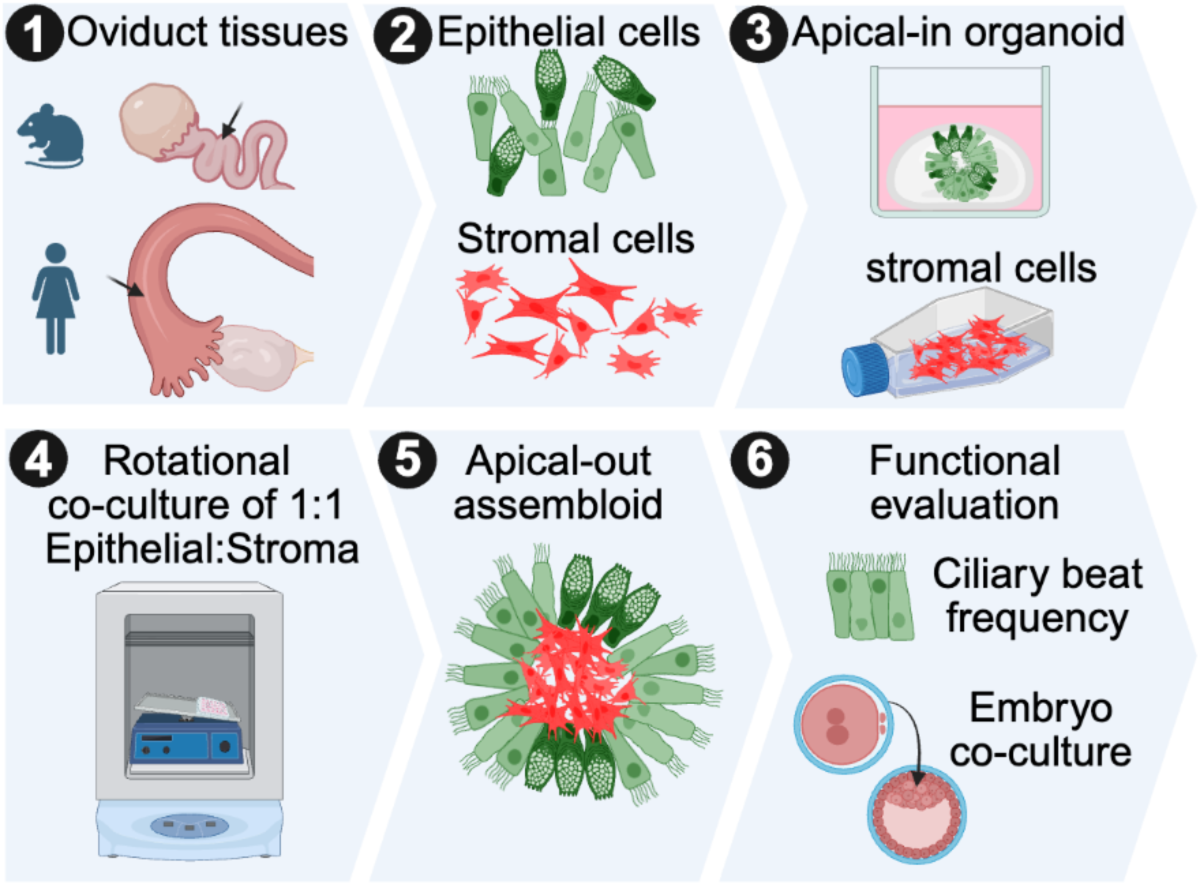

**Summary:** The protocol for generating mouse and human oviductal assembloids by combining epithelial and stromal cells for studying oviductal function in an *in vitro* setting.

## Introduction

The oviduct, or Fallopian tube in humans, is the site of fertilization and preimplantation embryonic development in mammalian species. Despite its essential role in fertility and early pregnancy establishment, current understanding has a limited ability to investigate the *in vivo* function of the oviduct during pregnancy establishment, primarily because of the difficulty accessing the tissues without removing the oviduct from the body cavity, especially in humans. In rabbits and sheep, the physiological function and secretory content of the oviduct have been studied using sedation, organ exteriorization, and oviductal cannulation [1–4]. Recent technological advancements have enabled studies of oviduct behavior using high imaging resolution in the presence of sperm and during preimplantation in *ex vivo* [5, 6] and *in vivo* environments [7–9]. Oviduct tissues collected directly from animals offer valuable insights, but *ex vivo* preparations can generally be maintained for short periods. Consequently, there is growing interest in developing a physiologically relevant 3D model for oviduct research.

Oviductal organoids are generated by allowing epithelial cells to self-organize into spherical structures within a solid basement membrane extract (BME) [10]. However, oviductal organoids in BME typically orient their apical surface toward the luminal core [11, 12], making it difficult to sample secretory content or to study interactions with gametes and embryos. Bovine oviductal epithelial spheroids cultured in media or synthetic oviductal fluid self-organized into apical-out spheroids, maintaining ciliary beating for at least 10 days [13, 14]. A recent study showed that apical-out human Fallopian tube organoids can be used to study interactions between ciliated cells and sperm *in vitro* [15]. However, epithelial-only organoids and spheroids do not fully reproduce stromal-epithelial signaling. Stromal cells are mesenchymal-derived fibroblasts that provide factors important for oviduct function, such as insulin-like growth factor-1 (IGF-1) [16]. Recent approaches combining epithelial and stromal components into self-organizing “assembloids” may more faithfully represent oviductal function and improve paracrine signaling [17, 18]. In conventional apical-in assembloids, however, microinjection remains necessary to access secretory fluids or examine cilia. We adapted our previously described mouse and human endometrial apical-out assembloid procedures [19, 20] to generate oviductal assembloids. Compared with conventional apical-in models, the apical-out feature provides direct access to the epithelial apical surface and surrounding culture media, facilitating studies of ciliary function and embryo co-culture. In this protocol, mouse or human oviductal organoids are designated as MOO or HOO, respectively, and the corresponding assembloids are designated MOA or HOA.

## Materials

Procedures for mouse studies were approved by the Animal Care and Use Committee (ACUC), and human tissue collection by the Institutional Review Board (IRB). In this study, *Wnt7a*^Cre/+^ mice [21] were bred with mTmG^f/f^ reporter mice [22] to selectively label epithelial cells with green fluorescent protein (GFP) in the reproductive tract, while the stroma and other cell types were labeled with red fluorescent protein (RFP). To reduce variability associated with cycle stage, oviducts were collected from adult (8-16 weeks old) female mice at the estrus stage. Human oviducts were collected from healthy subjects aged 18-45 who underwent salpingectomy for benign gynecological indications. Subjects had not used hormonal contraceptives for at least 3 months prior to the surgery. Most materials and reagents used in the following procedures are the same as those used in our previously published method for generating mouse and human endometrial assembloids [19], which are listed in Table of Materials 1. Before cell isolation, ensure all solutions and reagents are aseptically prepared and ready to use. Immunostaining of MOA/HOA can be performed as described for mouse and human endometrial assembloids [19] and is not repeated here.

### Step-by-step method details

#### 1. Mouse oviductal cell isolation

Day 1: Mouse tissue digestion

1.1. At 16:00h, collect oviducts from at least 3 mice in the estrus stage in 1× Ca^2+^/Mg^2+^- free Dulbecco’s phosphate-buffered saline (DPBS) in a 35-mm culture dish.

1.2. Wash the tissues in DPBS to remove blood and cellular debris.

1.3. Trim away the bursa and fat. Ensure that the uterine tissue surrounding the uterotubal junction is trimmed off (Figure 1A).

**Figure 1.**
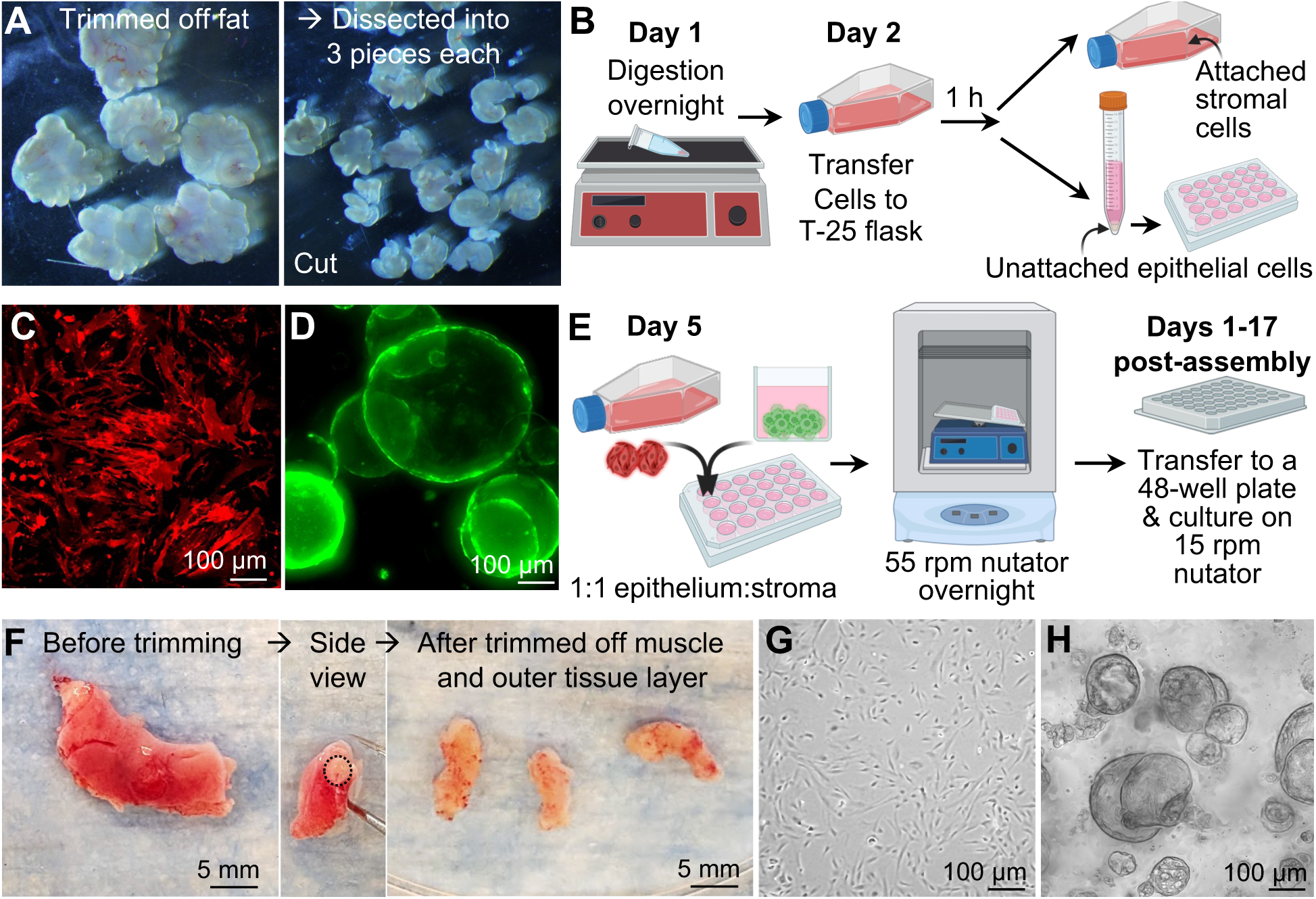
Generation of mouse and human oviductal organoids and assembloids. A. Mouse oviduct dissection after the fat was trimmed off and then dissected into 3 pieces for each oviduct. B. The tissue pieces were digested overnight on the orbital shaker and then transferred to the flask the following day. After 1 hour (h) of incubation, attached stromal cells are cultured, and unattached epithelial cells are centrifuged, resuspended, and plated in a 48-well plate. C. Red-fluorescent protein (RFP)-positive mouse stromal cells after 4 days of expansion. D. Green-fluorescent protein (GFP)-positive mouse epithelial cells organized into mouse oviduct organoids after 4 days of expansion. E. On day 5 after cell isolation, the stromal and epithelial organoids were combined at a 1:1 ratio and co-cultured in a rotational shaker (nutator) set at 55 rpm overnight. The following days 1-17, assembloids can be transferred to a 48-well plate and cultured on the nutator at 15 rpm. F. Human oviduct at the ampulla region before dissection. The tissue was then turned sideways to visualize the muscle and mesosalpinx area to be trimmed off (outside the circle). After trimming, only the epithelial and stromal layers are left and dissected into 5- to 7-mm pieces for digestion. G. Human stromal cells after 4 days of expansion. H. Human oviduct epithelial organoids after 4 days of expansion. B. and E. are created using BioRender.

1.4. Cut each oviduct into 3 pieces. Note: do not cut into overly small fragments, as small pieces can make it difficult to separate the epithelial sheet from the stroma after digestion.

1.5. Place tissues in 2 mL of cell dissociation solution in a 5-mL tube.

1.6. Incubate tissue overnight at 4°C, shaking gently on the orbital shaker with the speed set to 1 (Figure 1B). Note: Ensure all tools are cleaned with 70% ethanol, and that aseptic technique is maintained under the biological safety cabinet throughout the procedure to minimize contamination.

Day 2: Mouse stromal and epithelial cell isolation

1.7. At 09:00h, collect the tube from the orbital shaker. Add 1 mL DMEM/F12 supplemented with 10% fetal bovine serum (DMEM/F12+10%FBS) into the tube to stop the enzymatic reaction.

Note: Keep all media and cell suspensions on ice when tissues are not currently in use unless otherwise stated.

1.8. To separate the epithelial cell sheet and stromal cell layer from the connective tissue, pipette up and down 20 times using P1000.

1.9. Allow undigested tissue fragments to settle for 30 sec – 1 min.

1.10. Aspirate and transfer supernatant in a labeled 15-mL conical tube containing cell suspension.

1.11. Add 1 mL DMEM/F12+10% FBS to the undigested tissue fragments and pipette up and down 20 times using a P200, forcing tissue segments through the orifice.

1.12. Repeat steps 1.9-1.11, adding supernatant to your 15-mL tube, until the oviduct pieces no longer get stuck in the pipette tip and the supernatant is no longer cloudy (∼ 3 times).

1.13. Centrifuge at 300 × *g* for 5 min at 4°C.

1.14. Aspirate and discard the supernatant; save the cell pellet.

1.15. Add 1 mL DMEM/F12+10% FBS and pipette up and down 20 times to resuspend the cell pellet.

1.16. Transfer 1mL single cell suspension to a new labeled 5-mL tube.

1.17. Add 2 mL DMEM/F12+10%FBS to a 15-mL tube, then rinse the tube wall thoroughly to collect any remaining cells at the bottom. Transfer to a 5-mL tube.

1.18. Centrifuge at 300 × *g* for 3 min at 4°C.

1.19. Discard supernatant and save the cell pellet; do not put back on ice.

1.20. Add 1 mL of warm (37°C) Stroma cell growth media into the 5-mL centrifuge tube containing the cell pellet, then gently pipette up and down 20 times to resuspend the pellet.

1.21. Transfer all cell suspension into the T-25 flask containing 3 mL of Stroma cell growth media.

1.22. Add 2 mL Stroma cell growth media to the 5-mL tube, making sure to dispense media down the sides of the tube to rinse any remaining cells to the bottom, then transfer the cell suspension to a T-25 flask.

1.23. Rinse the 5-mL tube one more time with 1 mL of Stroma cell growth media and transfer to the T-25 flask, bringing the total media volume in the flask to 7 mL (Figure 1B).

1.24. Incubate the T-25 flask in a humidified 5% CO_2_ incubator at 37°C for 1 hr.

1.25. After 1 hr, examine the flask under the microscope to ensure that stromal cells are attached to the bottom of the flask. If not, wait for 15-30 min.

1.26. Tilt the flask and collect ∼6.5 mL of media containing unattached epithelial cells and transfer into a 15-mL conical tube on ice (Figure 1B).

1.27. Immediately and gently add 7 mL of prewarmed Stroma cell growth media to the side of the T-25 flask to avoid dislodging newly attached stromal cells, and continue the stromal cell culture in a humidified 5% CO_2_ incubator at 37°C.

1.28. Centrifuge the 15-mL conical tube containing epithelial cells at 300×*g* for 5 min.

1.29. Discard supernatant and resuspend the cell pellet in 1 mL cold Base organoid media.

1.30. Aspirate the single cell suspension and transfer to a new labeled 5-mL tube.

1.31. Add 2 mL Base organoid media to a 15-mL tube, making sure to dispense media down the sides of the tube to rinse any remaining cells to the bottom, then transfer to a 5-mL tube.

1.32. Centrifuge at 300×*g* for 5 min at 4°C.

1.33. Discard supernatant and resuspend the cell pellet in 500 μL Base organoid media.

1.34. Transfer the cell suspension to a new labeled 1.5-mL microcentrifuge tube.

1.35. Add 1 mL Base organoid media to a 5-mL tube, making sure to dispense media down the sides of the tube to rinse any remaining cells to the bottom, then transfer to a 1.5-mL tube.

1.36. Centrifuge at 300 × *g* for 5 min at 4°C.

1.37. Discard the supernatant and resuspend the cell pellet in 150 μL of Cultrex basement membrane extract (Cultrex BME) (add 50 μL more for each additional mouse used as a starting material).

Note: All manipulations with the Cultrex must be performed on ice to maintain a liquid state unless otherwise stated. Be cautious while pipetting, not to release air bubbles into the gel.

1.38. In a new 12-well plate, pipette four 15-μL droplets of resuspended cell-Cultrex mixture into each well (Figure 1B).

1.39. Cover the plate and gently invert it to create hanging drops. This will keep the cells from settling to the bottom while the Cultrex sets up.

1.40. Incubate in a humidified 5% CO_2_ incubator at 37°C for 15 min to polymerize the Cultrex.

1.41. Return the plate to the upright position and add 1 mL 37°C Mouse organoid expansion media to each well by dispensing down the side of the well. Do not directly spray the Cultrex droplets. Forceful pipetting may cause Cultrex to dissociate from the plate. If Cultrex is not well-polymerized, it will dissolve in the cell culture medium.

Note: The stromal cells are now cultured in the T-25 flask, and the epithelial cells are embedded in Cultrex in the 12-well plate.

#### 2. Preparation of stromal cells and mouse oviductal organoids (MOO) to generate mouse oviduct assembloids (MOA)

Day 5: Protocol 2 below is performed on the same day for both stromal cells and epithelial organoids. Cells may take 3-5 days to expand after plating. Perform the following steps on days 5-7.

##### Mouse stromal cells

2.1. On day 5, the stromal cells should be ∼80% confluent in the T-25 flask. The stromal cells should be RFP+ and exhibit a spindle-like morphology (Figure 1C). Aspirate and discard the media.

2.2. Wash twice with 3 mL prewarmed DPBS. Discard the DPBS after each wash.

2.3. Add 3 mL of TrypLE Express (room temperature).

2.4. Incubate at 37°C for 7 min. Cells will start to detach after 3-5 min.

2.5. Add 1 mL cold Stroma cell growth media to neutralize the enzyme activity and pipette up and down 10 times to break up the stromal sheets lifted from the bottom of the flask.

2.6. Add 2 mL of cold Stroma cell growth media to the flask, pipette up and down 5 times, and transfer to a 15-mL tube on ice using a 5-mL serological pipette.

2.7. Wash the flask with another 2 mL of Stroma cell growth media and collect it into a labeled 15-mL tube.

2.8. Centrifuge 300 × *g* for 5 min at 4°C.

2.9. Aspirate and discard supernatant, reserving cell pellet.

2.10. Resuspend cells in Stroma cell growth media. Pipette up and down a few times.

2.11. Centrifuge 300 × *g* for 5 min. Aspirate and discard supernatant.

2.12. Resuspend the stromal cells in 1 mL 37°C Base organoid media.

2.13. Count cells and assess viability (11 μL cell suspension + 11 μL Trypan blue) using Countess automated cell counter. Only samples with at least 75% cell viability will be used to generate assembloids.

##### MOO

2.14. On the same day, MOO should expand to ∼200-500 μm in diameter of apical-in GFP+ cells (Figure 1D).

2.15. Aspirate and discard the Mouse organoid expansion media

2.16. Remove Cultrex by adding 1 mL ice-cold DPBS into the 12-well dish. Dispense media at the base of the Cultrex bubble to remove it from the bottom of the plate and then aspirate and dispense into a 5-mL tube.

2.17. Once all MOOs are collected, pipette up and down 20 times with a P200 to break up organoids.

2.18. Incubate on an orbital shaker for 45-60 min at 4°C.

2.19. Ensure that the Cultrex gel is now homogeneous and in a liquid state.

2.20. Centrifuge 300 × *g* for 5 min at 4°C.

2.21. Aspirate and discard the supernatant. A layer of cells is visible at the bottom, then a layer of gel, topped by DPBS.

2.22. Remove the DPBS and gel, but not the cells. If the gel has re-solidified, rinse with another 3 mL of cold DPBS and centrifuge again. Repeat this step until the gel layer disappears.

2.23. Add 3 mL cold Base organoid media to the cell pellet and resuspend by pipetting up and down 10 times.

2.24. Centrifuge 300×*g* for 5 min at 4°C.

2.25. Remove supernatant and resuspend in 1mL 37°C Base organoid media.

2.26. After resuspension, the spheroid structure of MOO should be broken apart into epithelial sheets and single cells.

2.27. Count cells (11 μL cell suspension + 11 μL Trypan blue) using Countess automated cell counter.

##### Generation of MOA

2.28. Seed stromal and epithelial cells at a 1:1 ratio (10,000 stroma:10,000 epithelial cells) in a low-attachment surface 96-well plate. Ensure that each well contains a final volume of 120 μL of Base organoid media (Figure 1E).

2.29. Incubate on a nutator set to 55 rpm at 37°C, 5% CO_2_ incubator overnight.

2.30. Four days after MOA assembly (called day 4 MOA), MOA will have characteristics of apical-out epithelial cells surrounding the stromal core. MOA can now be transferred to a 48-well low-attachment plate. In general, ∼30 MOAs (or ∼75% of the total number of assembloids plated) are formed after 4 days using this cell ratio.

2.31. MOA can continue to be cultured for up to 17 days post-assembly on the nutator at 15 rpm in a humidified 5% CO_2_ incubator at 37°C (Figure 1E). Media are changed twice a week.

2.32. Use the following “gravity-based” method to transfer the MOA. Set the pipetter to ∼30 µL fitted with the wide-bore 200-μL tip. Pipette up and down, and examine the liquid in the pipette for small visible white dots of MOAs. If they are not visible, repeat pipetting and re-examine.

Note: When pipetting, aim to aspirate the liquid from the bottom of the plate, which generally contains MOA. The use of a 10× magnifying glass with a stand may facilitate identification of the MOA while working in the biosafety cabinet.

2.33. Once the MOA dots are aspirated into the tip of the wide-bore pipette tip, hold the pipettor vertically to let the MOAs “settle/fall” to the bottom of the pipette tip using gravity.

2.34. Lightly touch the bottom of the tip (containing MOAs) to the top of the media in the new well.

2.35. Wide-bore pipette tips can be used to handle MOA during culture, fixing, washing, and staining.

#### 3. Human oviductal cell isolation

Day 1: Human tissue digestion

3.1. Collect human oviduct tissues in RPMI+10% FBS with 1% antibiotic-antimycotic and keep on ice until arrival at the laboratory

3.2. Use sanitized forceps and scissors to handle the tissue during this stage

3.3. Trim away as much muscle and mesosalpinx tissue (the surrounding connective tissue around the stroma and epithelial cell layers) as possible without losing epithelial or stromal cells (Figure 1F, inside the circle).

3.4. Wash tissues in DPBS a few times to remove blood and cell debris.

3.5. Mince tissue into ∼0.5-mm pieces in DPBS and transfer to a 15-mL conical tube.

3.6. Place the 15-mL tube on an orbital rocker at 10 rpm for 5 min at room temperature.

3.7. Centrifuge at 100 × *g* at 4°C for 1 min.

3.8. Aspirate and discard supernatant.

3.9. Add 7 mL DPBS and repeat steps 3.6-3.8 until no visible red blood cells remain.

3.10. Add 4 mL of cell dissociation solution to the tissue in the 15-mL conical tube.

3.11. Incubate the tissues overnight at 4°C on an orbital shaker as stated above for mouse cell isolation protocols.

Day 2: Human stromal and epithelial cell isolation

3.12. At 09:00h, add 5 mL of DMEM/F12+10 % FBS to inactivate enzymatic activity.

3.13. Pipette up/down using a wide-bore P1000 pipette tip, allowing the tissue to settle at the bottom of the tube

3.14. Collect the top portion of the cell suspension (∼7.5 mL) and filter supernatant using a 40-μm cell strainer in a 50-mL conical tube.

3.15. Repeat steps 3.13-3.15 using a standard P1000 until the media no longer becomes cloudy.

3.16. Centrifuge at 300 × *g* at 4°C for 5 min.

3.17. Remove and discard the supernatant.

3.18. Add 1 mL DMEM/F12+10% FBS to resuspend the cell pellet and transfer to a labeled 15-mL tube.

3.19. Dispense 10 mL DMEM/F12+10% FBS media into the 50-mL tube to rinse. Transfer to a 15-mL tube.

3.20. Centrifuge at 300 × *g* at 4°C for 5 min.

3.21. Remove and discard supernatant.

3.22. Add 1 mL DMEM/F12+10% FBS to resuspend the cell pellet and transfer to a 5-mL tube.

3.23. Rinse the 15-mL tube with 3 mL of DMEM/F12+10%FBS and transfer to a 5-mL tube.

3.24. Centrifuge at 300 × *g* at 4°C for 5 min.

3.25. Do not put back on ice; aspirate and discard supernatant, and resuspend the pellet with 1 mL of prewarmed Stroma cell growth media

3.26. Transfer the resuspended cells into a T-25 flask containing 3 mL Stromal cell growth media.

3.27. Wash any potential leftover cells in the 5-mL conical tube with 2 mL Stromal cell growth media and transfer the resuspension to the T-25 flask

3.28. Repeat step 3.27 with 1 mL of Stroma cell growth media.

3.29. Total Cell-Stroma cell growth media volume is now 7 mL in the T-25 flask.

3.30. Incubate the cells in the flask for 30 min in a humidified 5% CO_2_ incubator at 37°C.

3.31. After 30 min, aspirate and transfer the media containing unattached cells to a new T-25 flask.

3.32. Add 7 mL of fresh Stroma cell growth media to the 1^st^ original T-25 flask and incubate both in a humidified 37°C, 5% CO_2_ incubator.

3.33. After an additional 45 min, collect media containing unattached epithelial cells from the 2^nd^ flask into a 15-mL conical tube.

3.34. Add 7 mL prewarmed Stroma cell growth media to the 2^nd^ T-25 flask and incubate both flasks containing stromal cells in a humidified 5% CO_2_ incubator at 37°C.

3.35. Proceed to follow steps 1.28-1.41, resuspend epithelial cells in ∼400 μL of thawed Cultrex, and use Human organoid expansion media, instead of Mouse organoid expansion media.

#### 4. Generation of human oviductal assembloids

Day 7: Protocol 4 below is performed on the same day. Cells may take 5-7 days to expand after plating. Perform the following protocol on days 7-9.

#### Human stromal cells

4.1. Once the stromal cells reach ∼80% confluency (Figure 1G), aspirate and discard the media.

4.2. Wash twice with 3mL prewarmed DPBS.

4.3. Add 3 mL of TrypLE Express into both flasks.

4.4. Incubate at 37°C, 5% CO_2_ incubator for 7 min.

4.5. Add 1 mL of Stroma cell growth media to neutralize enzymatic activity to each flask.

4.6. Mix well by pipetting up and down a few times using a serological pipette.

4.7. Add 2 mL to each flask and mix well.

4.8. Transfer cell suspension from both flasks to a single 15-mL conical tube.

4.9. Wash flasks to collect as many cells as possible with another 1 mL Stroma cell growth media and combine into the 15-mL conical tube. A total volume of collected cell+media is ∼14 mL.

4.10. Centrifuge at 300 × *g* at 4°C for 5 min.

4.11. Discard the supernatant and resuspend the pellet in 1 mL of Stroma cell growth media to wash. Transfer the single cell suspension to a 5-mL tube.

4.12. Wash the 15-mL tube with 3 mL Stroma cell growth media and transfer to a 5-mL tube.

4.13. Centrifuge at 300 × *g* at 4°C for 5 min.

4.14. Discard the supernatant.

4.15. Resuspend the pellet in 2 mL prewarmed Base organoid media.

4.16. Count cells and assess viability (11 μL cell suspension + 11 μL Trypan blue) using Countess automated cell counter.

#### HOO

4.17. On the same day, HOO should expand to ∼100-300 μm (Figure 1H).

4.18. Add ice-cold DPBS to dislodge Cultrex droplets. Transfer organoid suspension to a 5-mL tube.

4.19. Put on the orbital shaker and incubate for 60 min at 4°C.

4.20. Centrifuge at 300 × *g* at 4°C for 5 min.

4.21. Proceed to follow steps 2.19-2.23.

4.22. Remove supernatant and resuspend in 2 mL 37°C Human organoid expansion media.

4.23. Count cells and assess viability (11 μL cell suspension + 11 μL Trypan blue) using Countess automated cell counter.

#### HOA

4.24. Seed stromal and epithelial cells at a 1:1 ratio (10,000 stroma:10,000 epithelial cells) in a low-attachment 96-well plate, to a total of 80 μL media/well (50:50 Human organoid expansion media:Base organoid media).

4.25. Incubate on a nutator at 55 rpm overnight at 37°C and 5% CO_2_ incubator.

4.26. Transfer HOA to a new low-attachment 96-well plate with 80 μL Base organoid media/well and reduce nutator rotation speed to 15 rpm.

4.27. After ∼4 days, transfer the HOA to a 48-well low-attachment plate in Base organoid media and continue to culture at 37°C and 5% CO_2_ incubator on 15 rpm nutator.

4.28. Follow steps 2.31 – 2.36 to handle and transfer the HOA, similar to the steps for the MOA. Examples of MOA and HOAs are illustrated in Figure 2.

**Figure 2.**
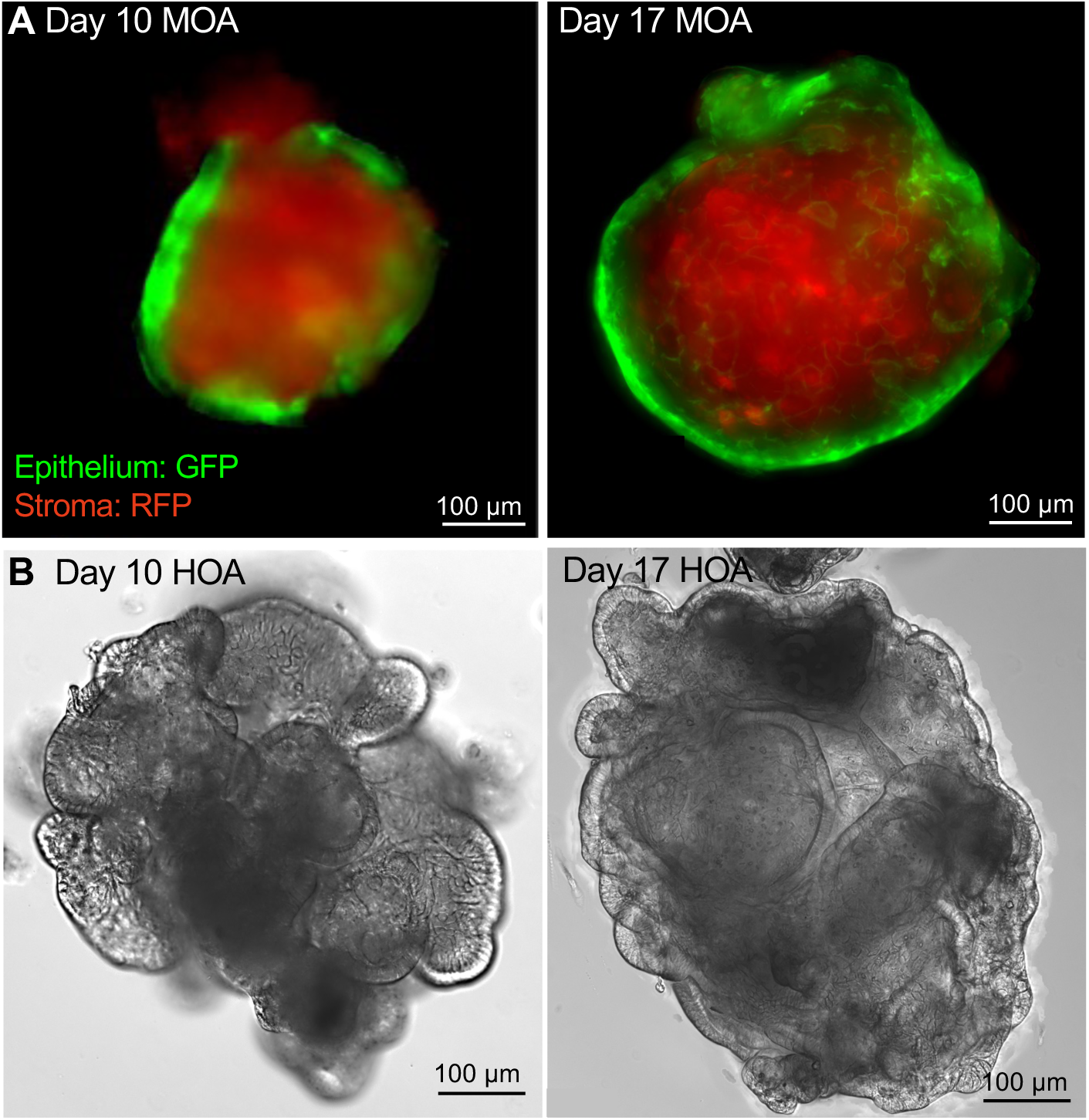
Morphology of mouse and human oviductal assembloids (MOA and HOA). A. Representative epifluorescence images of MOA generated from GFP+ epithelium and RFP+ stroma, derived from fluorescent reporter mice, at days 10 and 17 post-assembly. B. Bright field images of HOA showing distinct morphology and epithelial infoldings during culture at days 10 and 17 post-assembly.

#### 5. Assessment of the cellular identity and function of the assembloids

To confirm the cellular identity of assembloids, epithelial and stromal cells can be characterized using well-established cell markers. In both mouse and human oviducts, all epithelial cells are marked with E-cadherin, ciliated epithelial cells with acetylated α-tubulin (Ace α-Tub), secretory epithelial cells with oviductal glycoprotein (OVPG1), and stromal cells with Vimentin (VIM) (Figure 3). As ciliated epithelial cells are terminally differentiated, they often lose their beating function during *in vitro* culture. Ciliary beat frequency (CBF) can be another endpoint for assessing whether the oviductal epithelial cells retain their physiological function in assembloids following culture.

**Figure 3.**
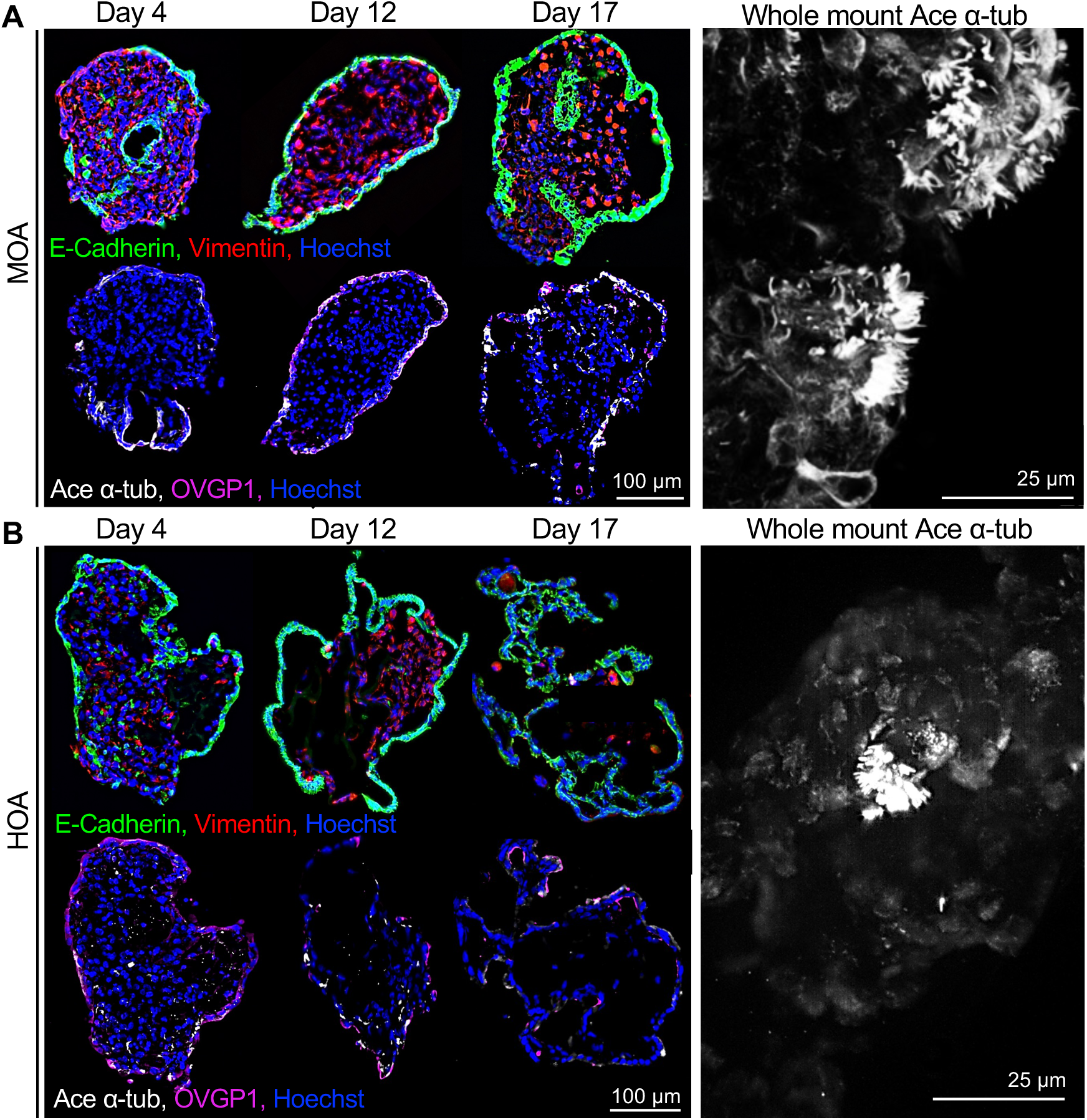
Cellular identity and apical-out polarity of mouse and human oviduct assembloids (MOA and HOA). Representative immunofluorescence images of epithelial and stromal compartments and ciliated and secretory epithelial cells in A. MOA and B. HOA collected at the indicated time points. Immunostaining identifies E-cadherin (green, epithelial cells), Vimentin (red, stromal cells), Ace α-Tub (white, ciliated epithelial cells), OVGP1 (magenta, secretory epithelial cells), and DNA (Hoechst, blue) in MOA and HOA. Whole-mount immunofluorescent images of Ace α-Tub immunostaining from day 10 MOA and day 8 HOA are shown in the right panels, demonstrating an apical-out localization of ciliated cells.

5.1. After tissue dissection, reserve a piece of mouse or human oviduct from the ampulla region for *ex vivo* controls. Keep the tissues in 37°C L15+1%FBS until imaging.

5.2. Slit the tissue lengthwise and transfer the cut tissue to the 35-mm glass-bottom dish containing 2-3 mL of L15+1%FBS media.

5.3. Use the slice anchor (“harp”) to pin down the tissue to the bottom of the dish, orienting the tissue so that the epithelial cell layer faces down. Make sure to wet the harp in L15+1%FBS before placing it on top of the tissue to help prevent tension from pushing the tissue out from under the harp when setting it down.

5.4. Let the tissue equilibrate in the heating chamber on the microscope for 30 min before imaging.

5.5. Using a 100× objective lens, record 10-second videos at a minimum of 60 frames per second where ciliated epithelial cells are visible.

5.6. Record at least 10 areas so that there are ∼ 500 ciliated cells for CBF quantification per sample.

5.7. With the MOA/HOA, transfer the assembloids using the “gravity” method into the 35-mm glass-bottom dish containing 200 μL of L15+1%FBS.

5.8. Record data from at least 10 assembloids/group.

5.9. To analyze the data, use the LAS X software with the “Stack Profile” tool in the “Quantify” tab to define a 2×2-pixel region of interest (ROI at the ciliary tip and measure changes in light intensity as the cilia move into and out of the ROI over time.

5.10. Move to the next ciliated cell or region until all data are collected for each sample.

5.11. Use a fast Fourier transform in R or AutoSignal to determine ciliary beat frequency in hertz.

#### 6. Co-culture of MOA and mouse embryos

To collect the zygotes, follow the protocol as described in the previous studies [23, 24].

6.1. The night before the culture, equilibrate the Potassium Simplex Optimized Media with Amino Acids (KSOM+AA) embryo culture media in the 37°C, 5% CO_2_ incubator by adding 150 µL of KSOM+AA to each 96-well plate (minimum of 5 wells).

6.2. Day 10 MOAs are used for the embryo co-culture experiment.

6.3. Using the “gravity” method described above, transfer the MOA into the 96-well plate containing 150 μL of KSOM+AA media.

6.4. Move the MOAs through 4 wells of 150 μL KSOM+AA to wash off the Base organoid media.

6.5. Note: Let the MOAs settle to the bottom of the well before transferring them to the next well.

6.6. Leave the MOAs in the 4^th^ well.

6.7. Using the capillary pipette, wash the zygotes through multiple microdrops of KSOM+AA and then add the zygotes to the 4^th^ well containing MOA at the ratio of 2:1 (zygotes:MOA), i.e., 10 zygotes: 5 MOAs per well.

6.8. Culture zygotes without MOA in the 5^th^ well as a negative control.

6.9. Record embryonic development daily through the blastocyst stage. Compare developmental timing and blastocyst formation between the MOA co-culture and KSOM+AA-only control groups.

### Representative results

When fluorescent reporter mice are used, typical MOAs contain lineage-specific cell populations of GFP+ epithelial cells surrounding an RFP+ stromal core (Figure 2A). In HOAs, invaginations of the epithelial cell layer should be visible by day 4 post-assembly, and persist through days 10-17 (Figure 2B). Unlike HOAs, MOAs did not exhibit excessive invaginations. Serial collection of assembloids can be used to assess cellular identity and function. In this study, MOA and HOA were collected on days 4, 12, and 17 post-assembly (Figure 3A-B). Immunostaining for E-cadherin and vimentin identified the outer epithelial layer and the stromal core, respectively. Acetylated α-Tub and OVGP1 identified ciliated and secretory epithelial cells, respectively. Whole-mount immunostaining for Ace α-Tub localized cilia to the external epithelial surface of MOA and HOA on days 10 and 8 post-assembly, respectively, demonstrating an apical-out epithelial orientation (Figure 3).

Ciliary beat frequency (CBF) provides a functional readout of the epithelial compartment. To provide a physiological benchmark, ampulla tissue was collected and analyzed as an *ex vivo* counterpart for mouse and human samples (Figure 4A and Supplementary Videos 1-2, respectively). CBF was quantified in MOA and HOA at days 10 and 8 post-assembly, respectively (Figure 4A and supplementary videos 3-4). Across randomly selected ROIs containing ciliated epithelial cells, CBF values ranged from 1 to 18 Hz in both *ex vivo* tissues and assembloids (Figure 4B).

**Figure 4.**
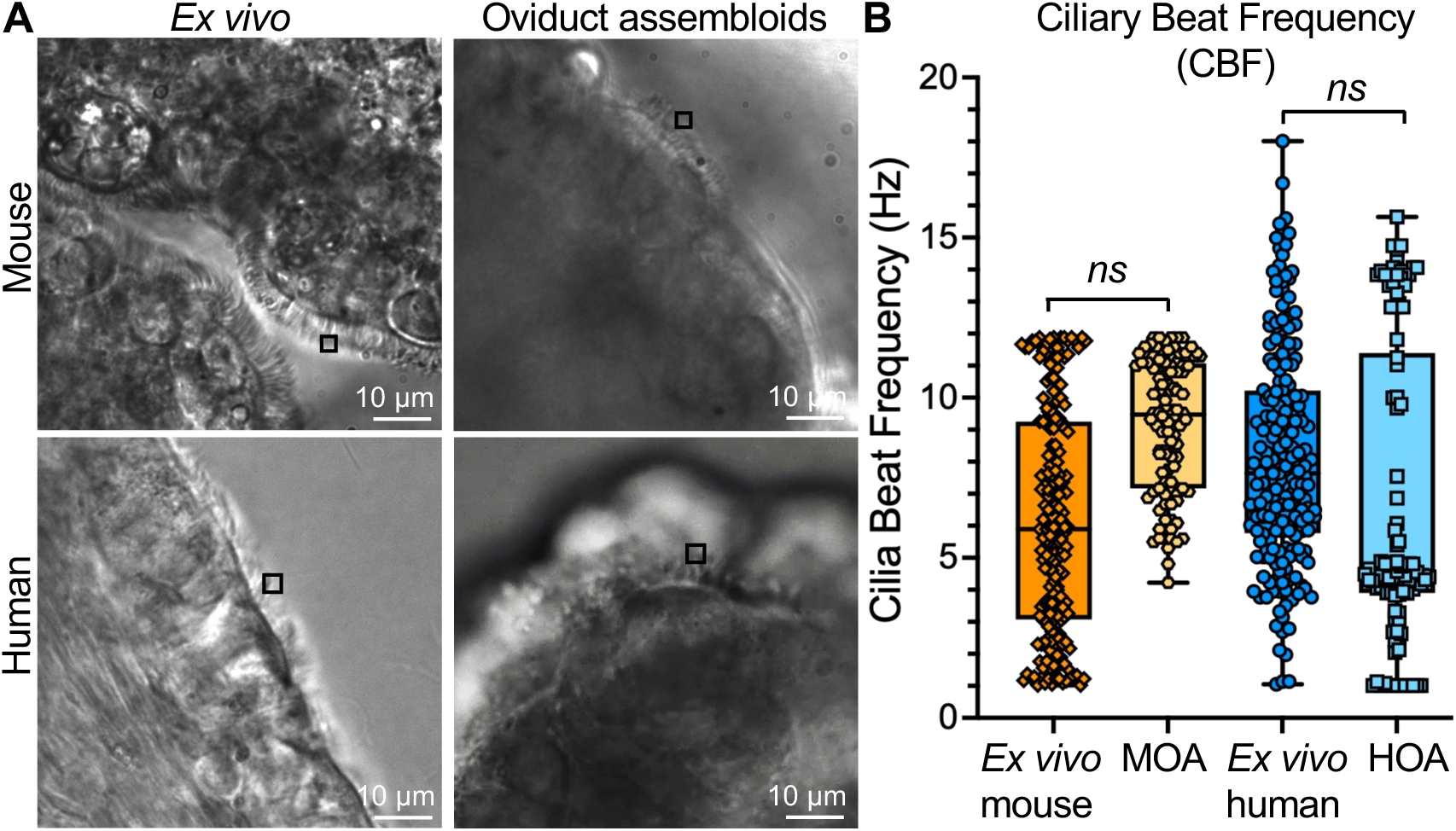
Ciliary beat frequency (CBF) in native oviduct tissue and oviductal assembloids. A. Representative still images of *ex vivo* mouse and human oviduct tissue, along with the corresponding MOA and HOA, used for ciliary imaging. B. Quantification of ciliary beat frequency in *ex vivo* tissues and corresponding assembloids. Each data point represents a single region of interest (ROI). Results were not statistically significant (*ns*); comparisons between corresponding species were made using an unpaired Student’s *t*-test.

A potential application of oviductal assembloids is the study of embryo-oviduct interactions and their use as a novel *in vitro* platform for evaluating early embryonic development. As a proof-of-concept experiment, day-10 MOAs were co-cultured with mouse zygotes. After 4 days of co-culture, zygotes cultured without MOA were predominantly either at the morula stage or underdeveloped (Figure 5A). The percentage of blastocysts developed in the media alone was significantly less than that collected from *in vivo* (Figure 5A-D). However, a higher proportion of embryos co-cultured with MOA progressed to the blastocyst stage, characterized by the formation of a blastocoel (Figure 5B). There was no statistical or morphological difference between embryos co-cultured with MOA and those collected from *in vivo* controls (Figure 5B-D). These observations support the feasibility of MOA-embryo co-culture for evaluating early embryonic development by studying the progression to the blastocyst stage.

**Figure 5.**
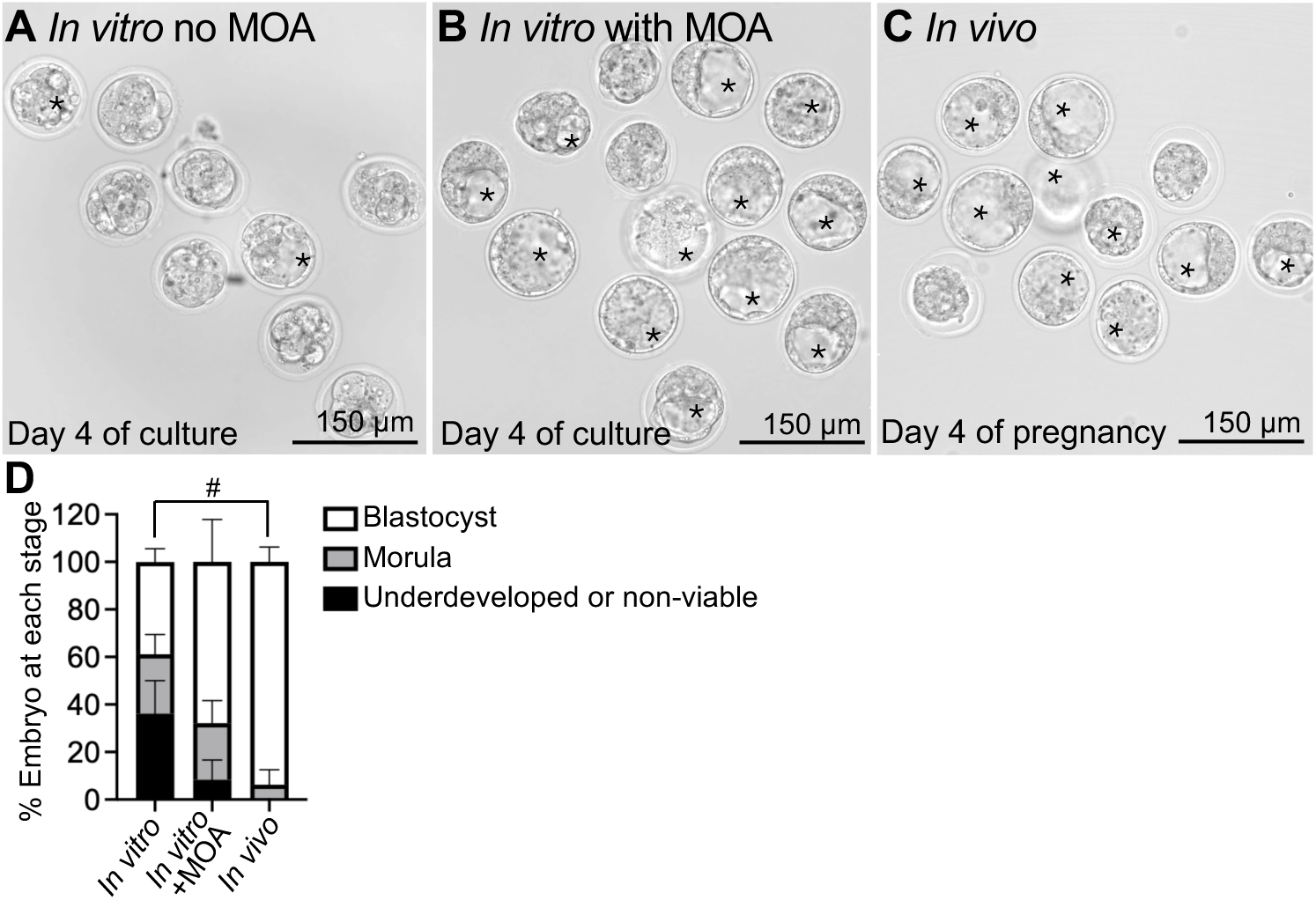
Mouse embryonic development with or without MOA compared to the *in vivo-*derived embryos. A. Representative embryos after 4 days of culture in KSOM+AA media without MOA. B. Embryos after 4 days of co-culture with KSOM+AA in the presence of MOA. C. Embryos flushed from the mouse uteri at day 4 of pregnancy. *, blastocyst-stage embryo. D. Embryonic developmental rate after 4 days of co-culture in the presence or absence of MOA compared to embryos flushed from mouse uteri on day 4 of pregnancy. ^#^*p*<0.05, %Blastocyst in *In vitro* group compared to *In vivo*. There was no statistical difference between the other groups. Two-way ANOVA with Tukey’s post-hoc test.

### Troubleshooting

Darkening of the stromal core may indicate necrosis. Potential contributing factors include reduced starting-cell viability, excessive assembloid size, inadequate medium exchange or agitation, poor nutrient diffusion, prolonged culture, or extended handling outside the incubator. Increasing cell input may enlarge the aggregate and limit diffusion. Instead, verify cell viability and counts, assembloid size, culture volume, nutator speed, and the time spent outside controlled temperature and CO_2_ conditions. Reduced or absent ciliary beating may reflect impaired epithelial cell function or mechanical damage during organoid recovery and assembloid transfer. Handle organoids and assembloids with wide-bore tips, minimize pipetting, and avoid direct contact with the external epithelial surface. Rotational movement of intact organoids attributable to ciliary activity can be recorded before assembly as an additional quality-control measure.

## Discussion

This protocol provides a framework for generating apical-out assembloids from mouse and human oviductal tissues. Mouse distal and proximal oviductal epithelial cells originate from distinct developmental lineages [25], and regional differences in organoid expansion have been reported in human and bovine tissues [12, 26]. Enrichment for ampullary tissue may increase expansion efficiency, but omitting the isthmus reduces representation of proximal oviduct biology. Therefore, the anatomical source of each preparation should be clearly documented, and regional cultures should be evaluated separately when investigating region-specific biology. In addition to the markers E-cadherin, vimentin, OVGP1, and Ace α-Tub described here, additional lineage-specific markers can be used to further characterize the cellular diversity in MOA and HOA, including ZO-1 and EPCAM for epithelial cells, PAX8 for secretory epithelial cells, FOXJ1 for ciliated epithelial cells, and COL1A1 for stromal cells [27, 28].

Assembloids should be allowed to self-organize and re-establish epithelial-stromal interaction for at least 4 days before downstream applications. The proof-of-concept MOA-embryo co-culture experiment suggests that MOA can support and may improve development through the blastocyst stage. As with other *in vitro* models, the oviductal assembloids model is also a reductionist by design and has inherent structural and technical limitations. MOA and HOA do not completely recapitulate the oviduct microenvironment, including smooth muscle, vasculature, immune cells, innervation, luminal flow, and region-specific anatomy [16, 23].

Outcomes are impacted by biological and technical variables, including animal cycle stage, human donor characteristics, tissue quality, and digestion efficiency, and batch-to-batch variation in extracellular matrix components. Additional studies should compare the hormonal responsiveness, transcriptomic profiles, secretory activity, metabolism, cellular composition, and long-term structural and functional stability of MOA and HOA with native oviduct tissues.

Oviductal assembloids may provide a platform for studying hormone responses, gamete-oviduct and embryo-oviduct interactions, and early cellular and molecular events associated with the development of high-grade serous carcinoma. However, the present human protocol relies on tissue collected during surgery for benign indications and has not yet been validated as a cancer model or drug-screening platform. Clinical translation to patient-derived malignant or precursor lesions would require confirmation that disease-associated genotypes and phenotypes are retained during organoids and assembloids generation and culture before therapeutic or personalized medicine applications are considered.

## Supplementary material

Supplementary videos 1-4 are available at BIOLRE online.

## Acknowledgement

The authors thank the Spencer/Kelleher lab staff for assistance in media and reagent preparation, Jeong/Kim laboratory for assistance with reporter mice, the Research Success Core at the University of Missouri (MU), Department of Obstetrics, Gynecology, and Women’s Health, for coordinating the procurement of human tissues from the MU Health Care Department of Pathology and Anatomical Sciences.

## Author contributions

KKS, AMK, and WW conceptualized the project. KKS, VA, AMK, and WW designed the experiments and protocols. MIH procured human tissues and maintained the IRB. KKS, VA, JAR, JM, AMK, and WW optimized the protocol. KKS, MAS, and MKS collected data. KKS and WW drafted the manuscript and figures. KKS, VA, MAS, MKS, JM, JAR, MIH, AMK, and WW edited the manuscript. AMK and WW provided funding for the project.

## Data availability

All the data were provided in the manuscript and the supplementary information.

## Table of Materials

| Materials | Company | Catalog number | Comments |
| --- | --- | --- | --- |
| ***Mouse models*** |  |  |  |
| *Wnt7a*<sup>Cre/+</sup> mice | Jackson Laboratory | 036637-JAX | Tg(Wnt7a-EGFP/cre)#Bhr/Mmjax |
| mT/mG<sup>f/f</sup> reporter mice | Jackson Laboratory | 007576 | *Gt(ROSA)26Sor<sup>tm4(ACTB-tdTomato,-EGFP)Luo/J</sup>* |
| ***Antibodies*** |  |  |  |
| Mouse monoclonal anti-E-Cadherin | BD Biosciences | 610182, RRID: AB_397581 | Immunofluorescent (IF), 1:1000 |
| Rabbit monoclonal anti-Vimentin | Cell Signaling | 5741S, RRID: AB_10695459 | IF, 1:1000 |
| Mouse monoclonal anti-acetylated α-tubulin | Millipore Sigma | T6793, RRID: AB_477585 | IF, 1:16000 |
| Rabbit polyclonal anti-oviductal glycoprotein 1 (OVGP1) | Invitrogen | PA520580, RRID: AB_11157467 | IF, 1:500 |
| Alexa Fluor 488, goat anti-mouse IgG | Jackson ImmunoResearch | 115545146, RRID: AB_2307324 | IF, 1:500 |
| Alexa Fluor 594, goat anti-rabbit IgG | Jackson ImmunoResearch | 111-585-144, RRID: AB_2307325 | IF, 1:500 |
| ***Chemicals, peptides, and recombinant proteins*** |  |  |  |
| Advanced DMEM/F-12 | Gibco | 12634028 |  |
| Leibovitz's L-15 medium | Gibco | 41300-039 |  |
| B27 Supplement | Gibco | 12587010 |  |
| Insulin-transferrin-selenium (ITS) | Gibco | 41400045 |  |
| Primocin | InvivoGen | NC9141851 |  |
| Glutamax | Gibco | 35050061 |  |
| A83-01 | BioGems | 9094360 |  |
| Murine EGF | Biotechne | 2028-EG-200 |  |
| Murine FGF-10 | Biotechne | 6224-FG-025/CF |  |
| Murine R-Spondin1 | Biotechne | 7150-RS-250/CF |  |
| Murine Noggin | Biotechne | 6997-NG-100/CF |  |
| Murine Wnt3a | Bio-Techne, R&D systems | 1324-WN-500/CF |  |
| Human EGF | Bio-Techne | AFL236-200 |  |
| Human FGF-10 | Bio-Techne | 345-FG/CF-MTO |  |
| Human R-Spondin1 | Bio-Techne | 4645-RS-01M/CF |  |
| Human Noggin | Bio-Techne | 6057-NG-100/CF |  |
| Human HGF | Bio-Techne | 294-HG-100/CF |  |
| N-acetyl-L-cysteine | Sigma | A9165 |  |
| Nicotinamide | BioGems | 9899208 |  |
| N-2 Supplenet 100X | Gibco | 17502001 |  |
| Y-27632 dihydrochloride | Sigma | Y0503 |  |
| Antibiotic-Antimycotic (100X) | Gibco | 15240062 |  |
| MgCl<sub>2</sub> (1 M) | Invitrogen | AM9530G |  |
| Cultrex BME | R&D Systems | 3433010-R1 |  |
| Dimethyl sulfoxide (DMSO) | Thermo Fisher Scientific | D128 |  |
| Fetal bovine serum (FBS) | Gibco | 16141079 |  |
| Collagenase V | Sigma | C9263 |  |
| DNase-I | Roche | 10104159001 |  |
| Liberase | Roche | 05401020001 |  |
| Hyaluronidase | Millipore Sigma | H4272 |  |
| TrypLE Express, no phenol red | Gibco | 12604021 |  |
| Dulbecco's Phosphate buffered saline (DPBS) | Millipore Sigma | D8537 |  |
| Borosilicate glass Pasteur pipette | VWR International | 89399-924 |  |
| KSOM + AA with D-Glucose | EmbryoMax | MR-106-D |  |
| Light Mineral Oil | Irvine Scientific | 9305 |  |
| Trypan Blue Solution | Gibco | 15250061 |  |
| Hoechst 33342 | Thermo Fisher Scientific | H3570 |  |
| Agarose Low Melting | Thermo Fisher Scientific | BP165-25 |  |
| Sodium citrate dihydrate | Thermo Fisher Scientific | BP327 |  |
| Paraformaldehyde | Thermo Fisher Scientific | 416785000 |  |
| Xylenes, histological grade | Sigma-Aldrich | 534056 |  |
| Ethanol, 200 proof (100%) | Decon Labs | 2705HC |  |
| Normal goat serum | MP Biomedicals | 08642921 |  |
| ProLong Diamond Antifade Mountant | Thermo Fisher Scientific | 36961 |  |
| Hematoxylin solution, Harris modified | Sigma-Aldrich | HHS32 |  |
| Calcium Sulfate, Drierite (Desiccant) | Alfa Aesar | 89751-P5 |  |
| Blue Ribbon Paraffin | Leica Biosystems | 3801360 |  |
| ART Wide Bore Filtered Pipette Tips 200 µL | Thermo Fisher Scientific | 2069G |  |
| ART Wide Bore Filtered Pipette Tips 1000 μL | Thermo Fisher Scientific | 2079G | |
| 12-Well Polystyrene Tissue Culture Plates | Cell Treat | 229112 |  |
| 48-well Ultra-Low Binding Culture Plate | ScienCell | 0343-4 |  |
| Nunclon Sphera 96U-Well Round Bottom Plates | Thermo Fisher Scientific | 174929 |  |
| 35-mm petri dish | Falcon | 351008 | For tissue collection |
| 35-mm glass-bottom dish with 100 micro-well | Cellvis | D35-10-0-N | For imaging ciliary beat frequency |
| 25 cm<sup>2</sup> (T-25) Polystyrene Tissue Culture Treated Flasks | CellTreat | 229331 |  |
| 1.5 mL polypropylene centrifugal tubes | Axygen | MCT150C |  |
| 5 mL polypropylene centrifuge tubes | Axygen | MCT500CS |  |
| 15 mL polypropylene centrifuge tubes | CellTreat | 229411 |  |
| 50 mL polypropylene centrifuge tubes | CellTreat | 229421 |  |
| 40 μm Cell strainer | Corning | 352340 | |
| 500 mL Filter System, 0.22 μm PES Filter | CellTreat | 229707 | |
| Tissue-Tek Cryomold | Sakura | 4565 |  |
| Tissue-Tek Mega Cassette System | Sakura | 4173 |  |
| FisherBrand Superfrost Plus Adhesion Microscope Slides | Thermo Fisher Scientific | 1255015 |  |
| SecureSeal™ Imaging Spacer | Electron Microscopy Sciences | 70327-9S |  |
| Microscope coverglass No.1 24×55 mm | Thermo Fisher Scientific | 12541033 |  |
| Microscope coverglass No.1 18 × 18 mm | Thermo Fisher Scientific | 12541012 |  |
| E-3063 tubing | Tygon | ACF00011 |  |
| **Equipment** |  |  |  |
| Micro Dissecting Scissors | Stoelting | 5213202P |  |
| High Precision Dissecting Micro Scissors, 4.75 in | Fisherbrand | 08-953-1B |  |
| High Precision Straight Slender Fine Point Tweezers/Forceps | Fisherbrand | 12-000-127 |  |
| Slice anchor for RC-42, 1.0 mm thread spacing, flat frame, stainless steel (Harp) | Warner Instruments | 64-1421 SHD-42/10 | To stabilize the tissue to the dish during ciliary imaging |
| Biological Safety Cabinet | Thermo Fisher Scientific | SN:300599231 |  |
| CO<sub>2</sub> incubator, HERAcell vios 250i | Thermo Fisher Scientific | SN42828487 |  |
| Countess Automated Cell Counter | Invitrogen | AMQAX2000 |  |
| Sorvall ST 16R bench centrifuge | Thermo Scientific | 75004381 |  |
| HistoCore Leica Pegasus Tissue Processor | Leica Biosystems | SN: 7958 |  |
| HistoCore Arcadia H Paraffin Embedding Station (heated module) | Leica Biosystems | SN: 7958 |  |
| HistoCore Arcadia C Paraffin Embedding Station (cold plate) | Leica Biosystems | SN: 8767 |  |
| Microtome | Leica Biosystem | RM2235 |  |
| Delcoking Chamber NxGen | Biocare Medical | DC121396 |  |
| CrestOptics CICERO spinning disk unit with the Leica DMI8 inverted microscope and 100x objective and 20x objective. Tokai Hit Stage Top Incubator | Leica Biosystem | SN: 678850 | For imaging cilia |
| Leica DM6 B upright microscope | Leica Biosystem | SN: 597425 |  |
| Leica DMI8 inverted microscope | Leica Biosystem | SN: 595981 |  |
| Stereo Microscope | Leica Biosystem | mz10f |  |
| Analog orbital shaker | Ohaus | 021061014 |  |
| Nutating Mixers - Variable Speed (Nutator) | Fisherbrand | 88-861-043 |  |
| Fisher Scientific 3D platform rotator | Thermo Fisher Scientific | 88861045 |  |
| Milli-Q IQ7000 Ultrapure Water Purification System | Millipore Sigma | ZIQ7000T0C |  |
| 2-in-1 LED 10× Magnifying Glass with Light and Stand | LANCOSC | n/a |  |

### Reagents, buffer and media components

#### Mouse/Human cell dissociation solution

**Reagents, buffer and media components** **Mouse/Human cell dissociation solution**
| Reagent | Stock concentration | Final concentration | Amount |
| --- | --- | --- | --- |
| RPMI | 1× | 1× | 1820 μL |
| Liberase | 5 mg/mL | 0.2 mg/mL | 80 μL |
| DNase-I | 10 mg/mL | 0.5 mg/mL | 100 μL |
| **Total** | **n/a** | **n/a** | **2000 μL** |

**Note:** Prepare fresh before each use.

#### Base organoid media

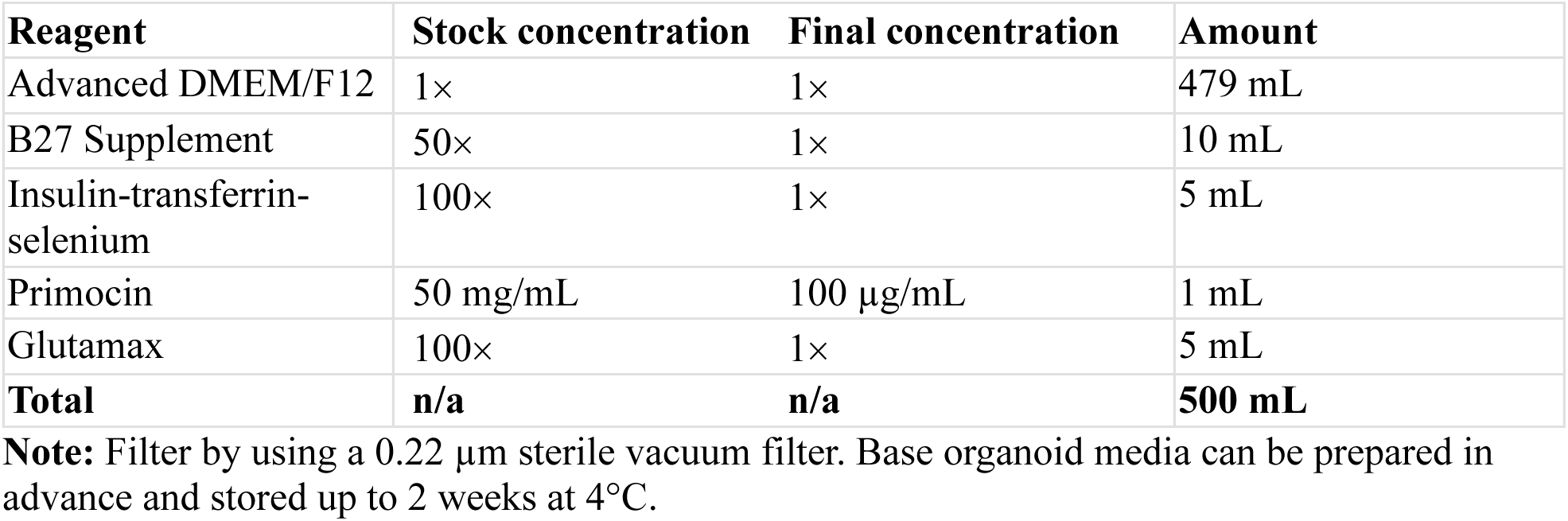

| Reagent | Stock concentration | Final concentration | Amount |
| --- | --- | --- | --- |
| Advanced DMEM/F12 | 1× | 1× | 479 mL |
| B27 Supplement | 50× | 1× | 10 mL |
| Insulin-transferrin-selenium | 100× | 1× | 5 mL |
| Primocin | 50 mg/mL | 100 μg/mL | 1 mL |
| Glutamax | 100× | 1× | 5 mL |
| **Total** | **n/a** | **n/a** | **500 mL** |

**Note:** Filter by using a 0.22 μm sterile vacuum filter. Base organoid media can be prepared in advance and stored up to 2 weeks at 4°C.

#### Mouse organoid expansion media

| Reagent | Stock concentration | Final concentration | Amount |
| --- | --- | --- | --- |
| Base organoid media | 1× | 1× | 196.4 mL |
| A83-01 | 5 mM | 500 nM | 20 μL |
| Murine EGF | 50 μg/mL | 50 ng/mL | 200 μL |
| Murine FGF-10 | 100 μg/mL | 100 ng/mL | 200 μL |
| Murine R-spondin1 | 200 μg/mL | 200 ng/mL | 200 μL |
| Murine Noggin | 100 μg/mL | 100 ng/mL | 200 μL |
| Murine Wnt3a | 50 μg/mL | 50 ng/mL | 200 μL |
| Nicotinamide | 1 M | 1 mM | 200 μL |
| N-2 Supplement | 100× | 1× | 2 mL |
| **Total** | **n/a** | **n/a** | **200 mL** |

**Note:** Mouse organoid expansion media can be prepared in advance and stored up to 2 weeks at 4°C.

#### Human organoid expansion media

| Reagent | Stock concentration | Final concentration | Amount |
| --- | --- | --- | --- |
| Base organoid media | 1× | 1× | 196.4 mL |
| A83-01 | 5 mM | 500 nM | 20 μL |
| Human EGF | 50 μg/mL | 5 ng/mL | 20 μL |
| Human FGF-10 | 100 µg/mL | 100 ng/mL | 200 µL |
| Human HGF | 50 µg/mL | 50 ng/mL | 200 µL |
| Human R-Spondin 1 | 500 µg/mL | 500 ng/mL | 200 µL |
| Human Noggin | 100 µg/mL | 100 ng/mL | 200 µL |
| Nicotinamide | 10 µM | 10 nM | 200 µL |
| N-acetyl-L-cysteine | 625 mM | 1.25 mM | 400 µL |
| N2 Supplement | 100× | 1× | 2 mL |
| **Total** | **n/a** | **n/a** | **200 mL** |

Note: Human organoid expansion media can be prepared in advance and stored up to 2 weeks at 4°C.

#### Stroma cell growth media

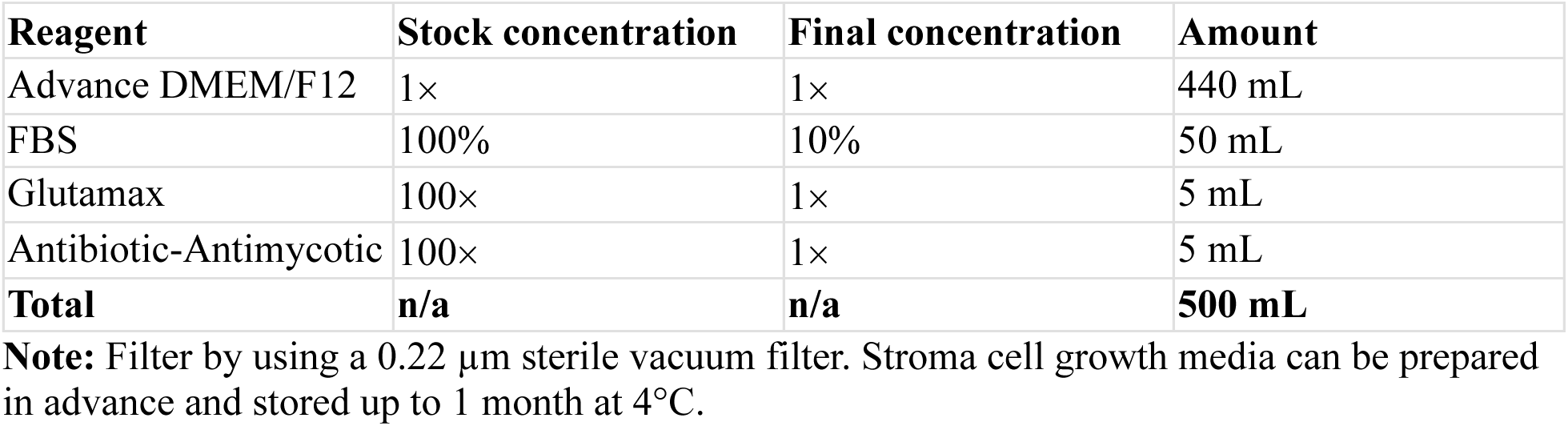

#### 4% Paraformaldehyde

| Reagent | Stock concentration | Final concentration | Amount |
| --- | --- | --- | --- |
| NaOH | 1× | 1× | 1.5 g |
| Paraformaldehyde (PFA) | n/a | 4% (wt/vol) | 4 g |
| DPBS | 10× | 1× (vol/vol) | 10 mL |
| ddH<sub>2</sub>O |  |  | ~90 mL |
| **Total** |  | **n/a** | **100 mL** |

Dissolve NaOH in 60 mL ddH<sub>2</sub>O. Add 4 g of PFA and stir for 20-30 minutes until fully dissolved. Add 10 mL of 10× DPBS and adjust pH to 7.2-7.4. Finally, add ddH<sub>2</sub>O to bring the total volume to 100 mL.

#### 1% Agarose low melting

| Reagent | Stock concentration | Final concentration | Amount |
| --- | --- | --- | --- |
| ddH<sub>2</sub>O | n/a | n/a | 100 mL |
| Agarose low melting | 100× | 1% (wt/vol) | 1 g |
| **Total** | **n/a** | **n/a** | **100 mL** |

Low melting point agarose solution can be prepared in advance and stored up to 4 weeks at room temperature.

#### Leibovitz’s L-15 media with fetal bovine serum (L15+1%FBS)

Leibovitz's L-15 media with fetal bovine serum (L15+1%FBS)
| Reagent | Stock concentration | Final concentration | Amount |
| --- | --- | --- | --- |
| Leibovitz's L-15 media | 1× | 1× | 1000 mL |
| FBS | 1× | 1× | 10mL |
| **Total** | **n/a** | **n/a** | **1100 mL** |

Preferably, hyaluronidase should be prepared fresh if L15+1%FBS is used. If not, DPBS can be used to replace L15+1%FBS and stored at 80°C for up to two months.

#### 0.1% Hyaluronidase

| Reagent | Stock concentration | Final concentration | Amount |
| --- | --- | --- | --- |
| Hyaluronidase | 1× | 1× | 2 mg |
| L15+1%FBS | 1× | 1× | 2 mL |
| **Total** | **n/a** | **n/a** | **2 mL** |

Preferably, hyaluronidase should be prepared fresh if L15+1%FBS is used. If not, PBS can be used to replace L15+1%FBS and stored at 80°C for up to two months.

#### Whole-mount blocking buffer

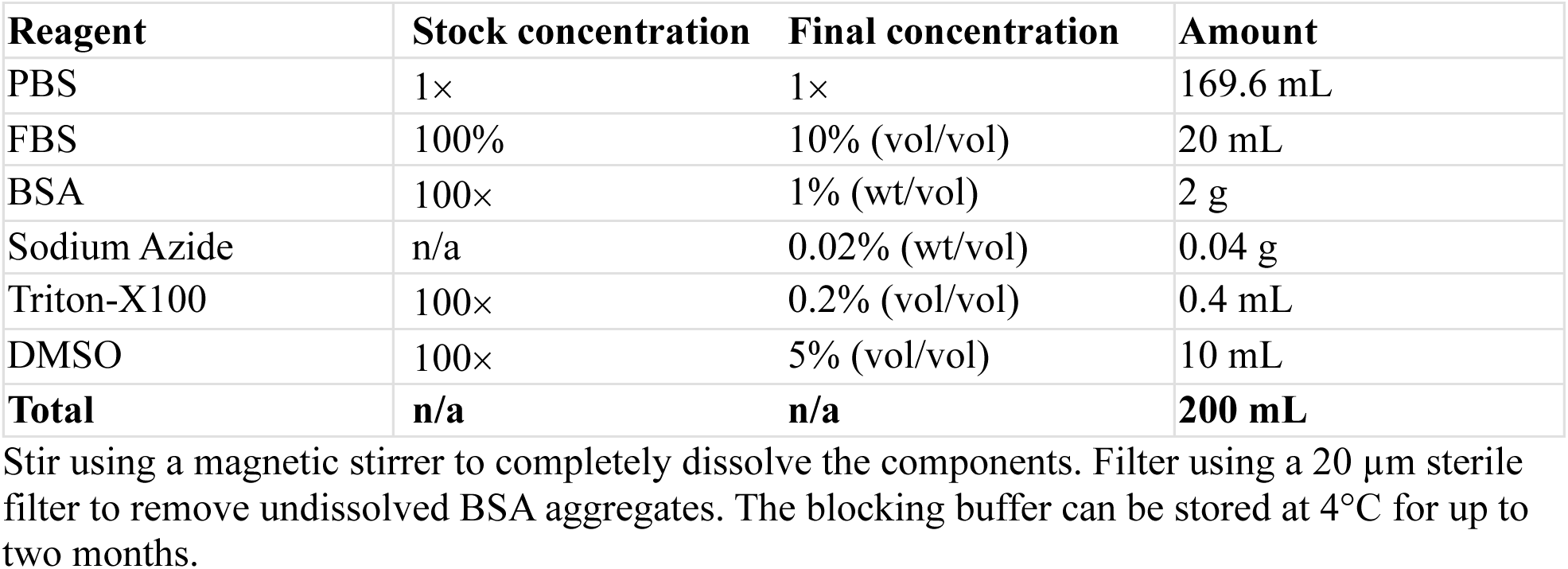

#### FLASH2 clearing reagent

| Reagent | Final concentration | Amount |
| --- | --- | --- |
| Urea | 25% (wt/vol) | 25 g |
| Borate (pH 7.0) | 200 mM (vol/vol) | 100 mL |
| Zwittergent | 8% (wt/vol) | 8 g |
| **Total** | **n/a** | **100 mL** |

**Note:** Urea dissolution in borate solution is an endothermic reaction that reduces Zwittergent solubility. It is critical to incubate the urea-borate solution on a heating plate to maintain room temperature. FLASH2 solution can be stored at 4°C for up to two months. Zwittergent precipitates at 4°C, so prewarm it to 60°C for approximately 3 min before each use.

#### Fructose-Glycerol Mounting Medium

| Reagent | Final concentration | Amount |
| --- | --- | --- |
| Glycerol | 60% (vol/vol) | 30 mL |
| Fructose | 2.5 M (wt/vol) | 22.52 g |
| ddH<sub>2</sub>O | n/a | 20 mL |
| **Total** | **n/a** | **50 mL** |

**Note:** Heat stir to completely dissolve fructose. Store at 4°C in a photo-resistant bottle. It can be used for 6 months.

## Notes

### Competing Interest Statement

The authors have declared no competing interest.

### Funding

- Eunice Kennedy Shriver National Institute of Child Health and Human Development: R01HD112315, R01HD097087

